# Aptamer based spatiotemporally controlled growth factor patterning for tunable local microvascular network formation in engineered tissues

**DOI:** 10.1101/2020.09.22.308619

**Authors:** Deepti Rana, Ajoy Kandar, Nasim Salehi-Nik, Ilyas Inci, Bart Koopman, Jeroen Rouwkema

**Affiliations:** Department of Biomechanical Engineering, Technical Medical Centre, Faculty of Engineering Technology, University of Twente, 7500 AE Enschede, The Netherlands; Izmir Democracy University, Vocational School of Health Services, Department of Dentistry Services, Dental Prosthetics Technology, Izmir, 35140, Turkey

**Keywords:** aptamers, vascular endothelial growth factor, patterning, programmable release, co-culture, vascularization

## Abstract

Spatiotemporally controlled growth factor availability is of crucial importance for achieving hierarchically organized vascular network formation within engineering tissues. Even though current growth factor delivery systems can provide sustained release and growth factor delivery on demand, they generally do not facilitate temporal control over the release rates and thus adaptation in accordance with the needs of growing engineered tissue. Additionally, with conventional growth factor loading methods, growth factors are often subjected to organic solvents or harsh conditions, leading to lower bioactivity and denaturation of the proteins. To overcome these limitations, this manuscript reports on the development of VEGF specific 5’ acrydite modified aptamer functionalized GelMA hydrogels. The covalently incorporated aptamers can selectively bind to proteins with high affinity and specificity, and can thus sequester the target protein from the surrounding environment. The manuscript shows that this not only provides temporal control over the growth factor release via complementary sequence hybridization, but also enables local control of microvascular network formation in 3D.

---

Achieving hierarchically organized vascular networks within engineered tissues is a long sought goal of tissue engineering and regenerative medicine.^[1]^ For this, it is important to understand and control the process of vascular network formation. The two major mechanisms that are responsible for natural vascular network formation are vasculogenesis (vessels form de novo) and angiogenesis (new vessels sprout off from pre-existing vessels). Both vasculogenesis and angiogenesis are driven by intrinsic genetic regulatory networks and the specific physicochemical properties of their immediate microenvironment, such as cell-cell interactions, cell-ECM interaction, and cell signaling via secreted or ECM-sequestered growth factors or cytokines.^[2]^ In order to mimic highly complex natural mechanisms like vasculogenesis or angiogenesis *in vitro*, there is an immediate need for the development of dynamic tissue engineered scaffolds that can be spatiotemporally controlled.

The spatial localization of growth factors such as vascular endothelial growth factor A (VEGF-A) plays a fundamental role in ensuring proper vascularization. VEGF-A has been shown to stimulate both vasculogenesis and angiogenesis. VEGF-A is a 45 kDa heterodimeric heparin-binding protein and exists in multiple isoforms that differ in their affinity for ECM binding.^[3][4]^ Studies in transgenic mice expressing only one of the major VEGF isoforms showed that their ability to bind ECM is critical for guiding proper vascular morphogenesis.^[3]^ Studies have also supported the idea of maintaining a balanced matrix-binding affinity of the VEGF rather than the presence of specific isoform is required for proper vascular morphogenesis.^[3]^ These observations altogether highlight the importance of spatially controlled bioavailability of growth factors within native tissue microenvironments. Additionally, multiple growth factors are required during different stages of vascularization. VEGF-A is for instance required to initiate endothelial cell recruitment and activation, and immature vasculature formation.^[3]^ At a later stage, platelet derived growth factor (PDGF-B) acts to recruit smooth muscle cells (SMCs) or pericytes for stabilization and remodeling of these immature vessels.^[5]^ Additionally, other growth factors like basic fibroblast growth factor (bFGF) and angiopoietins (Ang1 or Ang2) help in patterning and remodeling of vascular networks *in vivo*.^[6]^

Due to this complexity, a high level of spatiotemporal control on the bioavailability of multiple growth factors is needed for controlling vascularization within engineered tissues. Recent research has demonstrated the use of external stimuli such as light, enzymes, biomolecules, pH, or combinations of these, to achieve temporal control of growth factor availability.^[7–11]^ However, current growth factor delivering systems generally focus on the immobilization of growth factors via specialized linker proteins or peptides, such as matrix metalloproteinase-sensitive linkers, RGD based peptides or heparin binding peptides.^[12–16]^ Even though these approaches provide temporal control over growth factor delivery, there are limitations such as spatial selectivity and the inability to adapt release rates to the need of the developing engineered tissue.

To address aforementioned challenges, this study harnesses the unique properties of aptamers in achieving spatiotemporal control over growth factor availability for regulating vascularization within engineered tissues. Aptamers are single-stranded oligonucleotides selected from synthetic RNA/DNA libraries. They are small and structurally stable with low immunogenicity. Based on their sequence, these oligonucleotides form 3D conformations and bind to target molecules with high affinity and specificity.^[17,18]^ Additionally, aptamers show a high affinity to hybridize with complementary sequences, leading to a release of the target molecule.^[19]^ By exploiting the mechanisms of molecular recognition among aptamers, growth factor and its complementary sequence, growth factor delivery can be achieved by using complementary sequence as trigger molecules. Additionally, as the affinity of aptamers and their complementary sequences are easily tunable, the release rate can be made adaptable according to the needs of developing engineered tissues.

In order to incorporate aptamers within a bioactive hydrogel system, we synthesized aptamer-functionalized gelatin methacryloyl (GelMA) hydrogels using photo-crosslinking via free radical polymerization initiated by UV light exposure **(Figure 1A)**. We compared unmodified VEGF165 specific aptamer^[17,18]^ (hereafter referred as “control aptamer”) and 5’-acrydite modified aptamer (hereafter referred as “acrydite aptamer”) **(Figure S1, Table S1)**. Since the degree of methacrylation and the concentration of GelMA monomer is critical for acrydite aptamer incorporation as well as polymer network formation, GelMA with a medium methacrylation degree (∼60%) was used for synthesis **(Figure S2)**.

**Figure 1.**
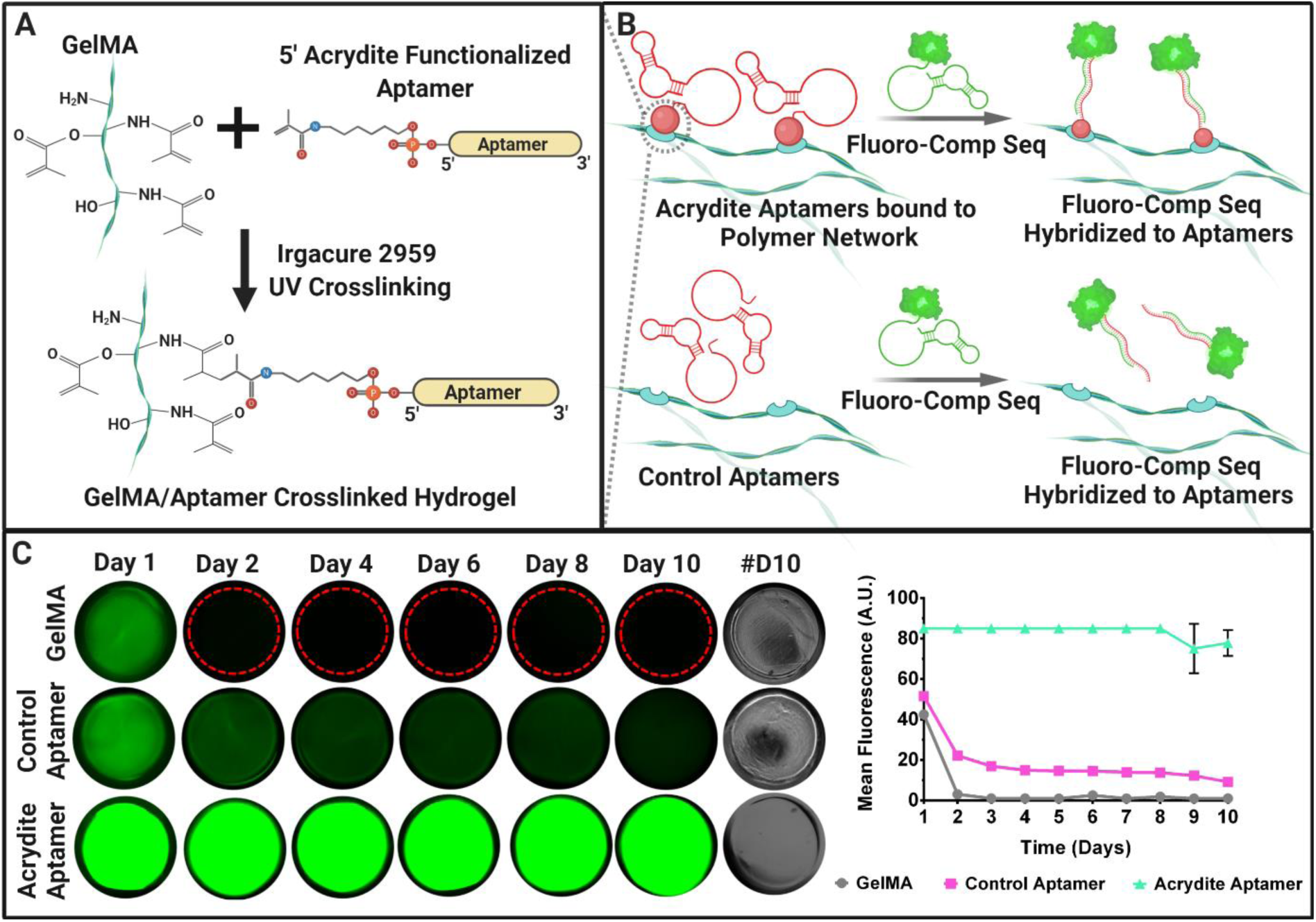
The concept of aptamer functionalized hydrogels and aptamer retention within the hydrogels. (A) Schematic representation of photo-crosslinked gelatin methacryloyl (GelMA) with 5’acrydite modified VEGF165 specific aptamer sequence, in the presence of photoinitiator (Irgacure 2959) and UV light. (B) Schematics explaining the concept of 5’acrydite modified aptamers being covalently linked into the GelMA network, whereas, due to the absence of 5’acrydite group (red circle), the control aptamers are diffusible within the hydrogel. Due to molecular recognition, upon addition of 5’-fluorescently labelled complementary sequence (Fluoro-Comp. Seq.) in the 5’acrydite functionalized hydrogels (Acrydite Aptamers) and control aptamer functionalized hydrogels (Control Aptamers), hybridization between the Fluor-Comp. Seq. and aptamers will occur. (C) The fluorescence microscopic images of the GelMA, Control Aptamer and Acrydite Aptamer hydrogels after 24 hrs incubation with Fluoro-Comp. Seq. at 37 °C. #D10 indicates the brightfield images of the hydrogels on day 10. The fluorescence intensity corresponding to the presence of Fluoro-Comp. Seq. indicates the presence of aptamers within the hydrogels at various time points. The red dotted line in GelMA hydrogels from day 2 onwards shows the hydrogel border. The mean fluorescence of all the samples over different time points have been shown in the graph. The experiment was performed with n=3 experimental replicates and values are represented as mean ± SD.

## Aptamer retention and molecular recognition with fluorescently labelled complementary sequence at physiological conditions

To verify the retention of VEGF specific acrydite- and control aptamers within the hydrogels, as well as their ability for complementary sequence recognition at physiological conditions, were performed. For this purpose, aptamer supplemented hydrogels were incubated with 5’-Alexa Fluor 488 fluorophore labelled complementary sequence for 24 hrs, followed by thorough washing and supernatant replacement with fresh DPBS after every 24 hrs until day 10 **(Figure 1B)**. The acrydite aptamer functionalized hydrogels displayed highest fluorescence (77 a.u.) which remained stable for 10 days of incubation at a physiological temperature (i.e., 37 C), confirming aptamer incorporation and complementary sequence recognition **(Figure 1C)**. On the contrary, the control aptamer-functionalized hydrogels and GelMA hydrogels showed lower fluorescence of 51 a.u. and 42 a.u., respectively after 24 hrs and their fluorescence decreased over time. In the case of control aptamers-functionalized hydrogels, the fluorescence decrease was gradual. In GelMA hydrogels fluorescence was no longer detectable after 48 hrs. These results indicate that control aptamers are physically entrapped and diffuse out of the hydrogels over time **(Figure 1B)**. Along with its retention capability, the results confirmed the high molecular recognition ability of the fluorescently labelled complementary sequence with the aptamers within the hydrogel matrices at physiological conditions, regardless of being acrydite functionalized or not.

## VEGF sequestration and its triggered release via complementary sequence hybridization

The functionality of aptamer-functionalized hydrogel in sequestering and regulating the release of VEGF, in the presence or absence of the complementary sequence, is determined primarily by the molecular recognition among these three molecules. In order to validate VEGF sequestration in the functionalized hydrogels and triggered release behavior via complementary sequence hybridization, aptamer-functionalized hydrogels (50 μl) were incubated with 10 ng of VEGF in 1 ml loading medium (0.1% BSA in DPBS) for 1 hr at 37 °C. Subsequently, samples were washed to remove unbound VEGF and supplemented with 1ml releasing medium (0.1% BSA in DPBS), with medium change after every 24 hr until day 10. GelMA hydrogels without aptamers were included as control. To examine the triggered release behavior, complementary sequence was added to releasing medium on day 4 and on day 9 **(Figure 2A, B)**. The mole ratio of aptamers to complementary sequence was fixed at 1:1. The concentration of VEGF in the releasing medium was determined using ELISA every 24 hrs.

**Figure 2.**
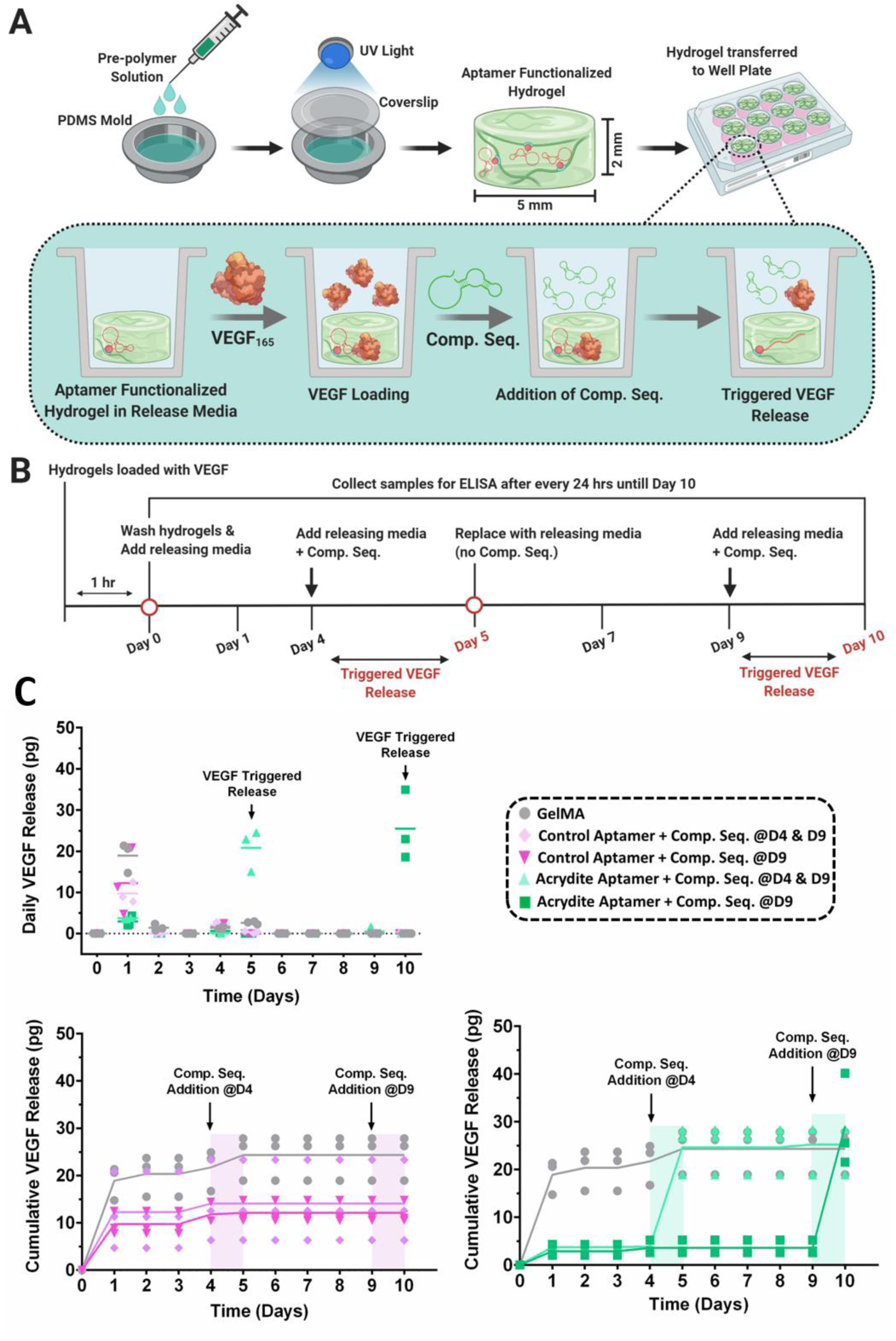
Spatiotemporally controlled VEGF release from the aptamer-functionalized hydrogels. (A) Schematic representation of the fabrication of aptamer-functionalized hydrogels, its ability to sequester VEGF from the release media via affinity interaction and on-demand triggered release of VEGF in the presence of complementary sequence (Comp. Seq.). (B) The complete timeline of the experiment. The aptamer concentration used was 25 μM and initial VEGF loading concentration was 10 ng. (C) The daily (up) and cumulative (down) VEGF release from control aptamer and acrydite aptamer functionalized hydrogels over 10 days in presence or absence of Comp. Seq. To trigger the first round of VEGF release on day5, the Comp. Seq. was added to specified hydrogels on day 4 for 24 hrs (as indicated in figure). Furthermore, to trigger the second round of VEGF release, the Comp. Seq. was added to all aptamer hydrogel samples on day9, except for control GelMA hydrogels. The mole ratio of Comp. Seq. to aptamers used was 1:1. The graphs are represented as mean with individual data points. The experiment was performed with three experimental replicates, n=3.

When examining the VEGF retention after 1 hr loading of the hydrogels **(Figure S3)**, aptamer-functionalized hydrogels showed higher VEGF retention (50% and 46% for control aptamer and acrydite aptamer respectively) compared to the GelMA hydrogels (33%). The increased retention in aptamer-functionalized hydrogels is likely due to the combined effect of diffusion and affinity binding. It should be noted that gelatin-based hydrogel matrices also show inherent growth factor sequestration properties due to their electrostatic interactions with oppositely charged growth factors like VEGF, which can explain the relatively high sequestration for the control GelMA hydrogels.^[20]^

Having confirmed the higher VEGF sequestration of aptamer-functionalized hydrogels after initial 1hr loading, the prolonged VEGF retention capability and triggered release behavior via complementary sequence hybridization was determined in physiological conditions **(Figure 2B)**. Control aptamer-functionalized and GelMA hydrogels showed a high burst release of VEGF on day 1 (10.99 pg and 18.89 pg respectively) and a near zero release from day 2 onwards. In contrast, acrydite aptamer-functionalized hydrogels showed a minimal initial burst release of 3.30 pg, indicating a good retention of the loaded VEGF. Upon the first round of triggered release with complementary sequence on day 4, the acrydite aptamer hydrogels showed a release of 20.81 pg on day 5, which was high compared to GelMA and control aptamer hydrogels (2.62 pg and 0.32 pg respectively). Upon the second triggered release on day 9, the acrydite aptamer and control aptamer hydrogels that were already triggered on day 4 showed no additional release. In contrast, the acrydite aptamer hydrogel that was triggered only on day 9 showed a high release of 25 pg on day 10 **(Figure 2B)**.

The acrydite aptamer hydrogels showed stable VEGF retention throughout the experiment and on-demand triggered release on day 5 and day 10 due to affinity based interactions. The cumulative VEGF release showed an overall release in the range of 25 – 29 pg in acrydite aptamer and GelMA hydrogels by day 10 **(Figure 2B)**, whereas the control aptamer hydrogels showed a cumulative release in the range of 12 – 14 pg **(Figure 2B)**. As the control aptamer is only physically entrapped within the polymer matrix, a portion of VEGF molecules diffusing out into the releasing medium is likely to be bound to the control aptamer. This correlates directly with the retention data of Figure 1, where the fluorescence was observed to reduce over time. To investigate whether VEGF molecules bound with control aptamers result in a lower signal of the ELISA assay, control aptamers solution (2.5 nmoles) in PBS supplemented with 10 ng VEGF was compared with PBS supplemented with 10 ng VEGF **(Figure S4)**. This resulted in a lower ELISA signal for control aptamer samples compared to PBS, confirming an effect of aptamer binding on ELISA sensitivity **(Figure S4)**.

It is important to note that in the presented approach, growth factor loading is decoupled from the hydrogel fabrication steps. This provides a clear advantage over conventional growth factor loading methods, where growth factors are often subjected to organic solvents or harsh conditions, leading to lower bioactivity and denaturation of the proteins.^[19,21]^ Additionally, target growth factors can be loaded at different time points after hydrogel fabrication, according to the need of the developing tissue. This provides great temporal control for tissue engineering applications.

## Aptamer Incorporation influences physicochemical properties

In order to assess whether aptamer incorporation affects the physicochemical properties of GelMA hydrogels, storage modulus and loss modulus were evaluated using rheological measurements **(Figure 3B, C)**. Control aptamer functionalized hydrogels showed an increase in storage modulus with increasing aptamer moles (∼390 Pa for 0.25 nmoles, ∼430 Pa for 2.5 nmoles and ∼509 Pa for 25 nmoles respectively) **(Figure 3B)**. Acrydite aptamer functionalized hydrogels on the other hand showed a storage modulus of ∼450 Pa, independent of changes in aptamer moles. The GelMA hydrogels exhibited a storage modulus of ∼410 Pa **(Figure 3C)**. While the exact reason for this difference between control aptamer and acrydite aptamer functionalized hydrogels is unclear, one possibility may be linked to the hydrophilic nature of aptamers resulting in additional swelling of the hydrogels, and thus in an increased storage modulus. This will be especially the case for control aptamers, which are only physically entrapped in the hydrogel network. Acrydite aptamers on the other hand are covalently incorporated into the hydrogel network, and are thus less freely available to induce swelling.

**Figure 3.**
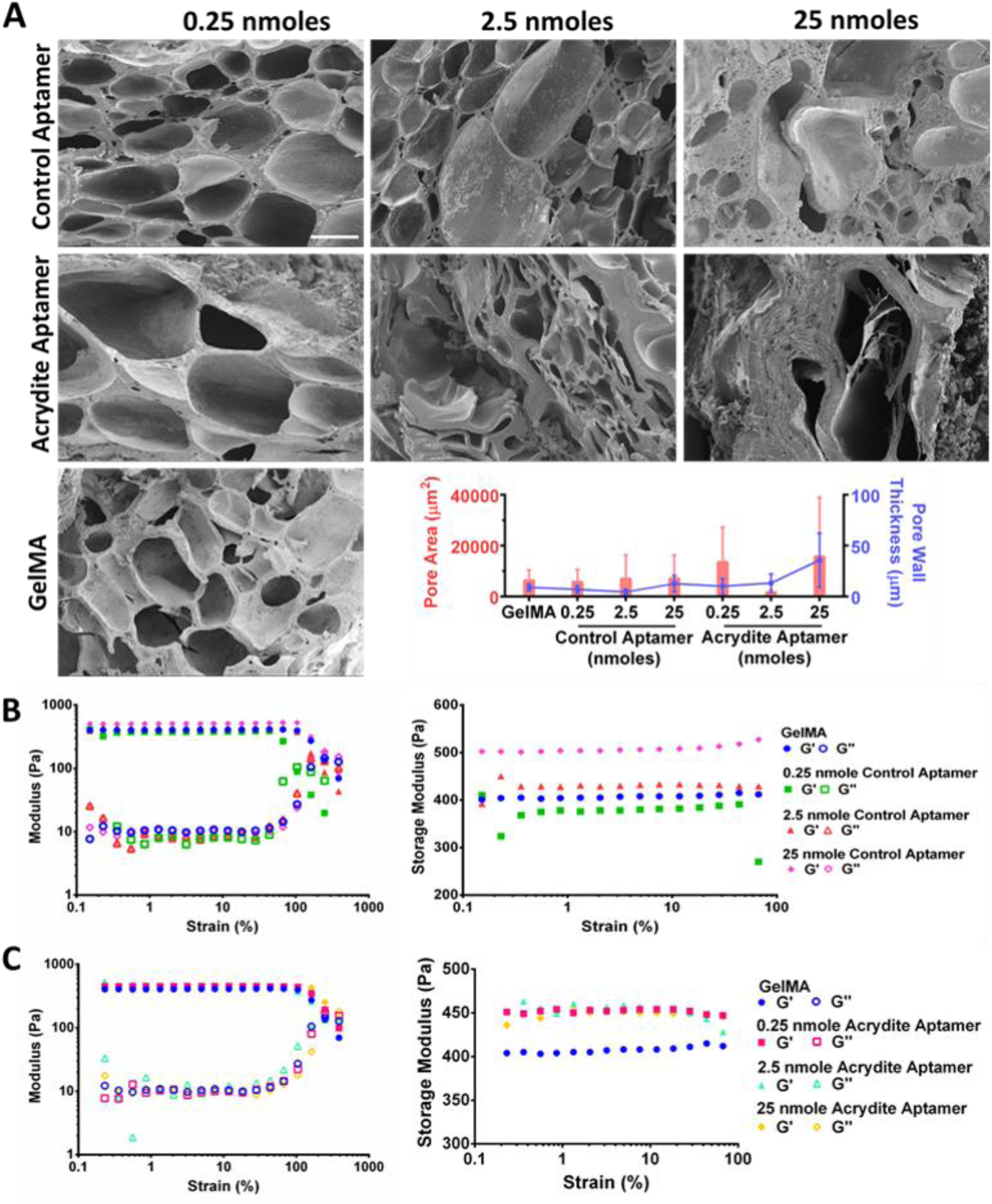
Physicochemical properties of aptamer functionalized hydrogels with different aptamer moles, namely 0.25 nmoles, 2.5 nmoles and 25 nmoles aptamers per sample. (A) SEM images of the cross-section of acrydite aptamer- and control aptamer-functionalized hydrogels with different aptamer concentrations. As a control, GelMA hydrogels without aptamers were used. Scale bar is 100 μm and magnification is 200x. Pore area and pore wall thickness of different hydrogel samples quantified by ImageJ using the representative SEM images. The data is represented as mean ± S.D. Rheological analysis of the (B) control aptamer-functionalized and (C) acrydite-functionalized hydrogel samples, along with control GelMA hydrogel samples. The storage modulus (filled symbols) and loss modulus (empty symbols) of the respective hydrogel samples at a fixed angular frequency of 1 rad/s is shown.

To further analyze the influence of aptamer incorporation on hydrogel pore properties, scanning electron microscopic (SEM) images of freeze-dried samples were examined. The results indicated similar pore properties among control aptamer functionalized hydrogels and GelMA hydrogels (**Figure 3A**). The average pore area for these hydrogels were found to be in the range of 6,000 to 6,900 μm^2^. In contrast, acrydite aptamer functionalized hydrogels showed mole dependent effect on their pore properties, where lower number of aptamer moles (0.25 nmoles) resulted in an increased average pore area of 13,320 μm^2^. Increase in aptamer moles resulted in densification of the hydrogel as shown by an increase in wall thickness and decrease in pore size for the 2.5 nmoles samples (∼13 μm and 1,462 μm^2^), and a collapse of pores as shown by a further increase of wall thickness and large remaining pores for the 25 nmoles samples (∼35 μm and 15,551 μm^2^). It should be noted that the hydrogel pore size as determined using SEM, does not represent the hydrogel ultrastructure of swollen hydrogels that is experienced by cells. Instead, the observed porosity is a result of the freezing and lyophilization process and represents the density of the swollen hydrogel samples.

## Cellular response to VEGF loaded aptamer-functionalized hydrogel in 3D culture conditions

We next demonstrated the biocompatibility of this system during VEGF sequestration and complementary sequence hybridization. Where previous studies have assessed the bioactivity of aptamer bound VEGF in an indirect way by culturing cells in medium conditioned with released VEGF,^[22,23]^ our aim was to investigate vascular network formation within the aptamer functionalized hydrogels. For this, pre-polymer solutions were mixed with human umbilical vein endothelial cells (HUVECs) and human mesenchymal stromal cells (MSCs) in a 1:1 ratio and subjected to photo-polymerization to yield cell-laden aptamer-functionalized hydrogels. These were incubated with 10 ng VEGF in 1ml of co-culture medium for 1 hr at 37 °C to enable VEGF sequestration. Subsequently, hydrogels were washed and medium was replaced with fresh co-culture medium. Cell viability was high (>90%) in acrydite functionalized hydrogels and GelMA controls on day 1 **(Figure S5)**. Cell viability remained high by day 5, independent of complementary sequence addition. In contrast, the control aptamer hydrogels showed a lower cell viability (>70%) on day 1. This increased significantly to >90% by day 5, independent of complementary sequence addition **(Figure S5)**. The initial lower viability in control aptamer samples remains unclear, but could be related to the cells preference for matrix bound growth factors in comparison to soluble ones.^[24]^ This cell viability data confirms the biocompatible nature of the system and shows no negative effect of VEGF sequestration as well as complementary sequence hybridization on 3D encapsulated cells.

In order to understand the effect of spatially patterned matrix bound growth factors on the surrounding cells, we designed bi-phasic cell-laden hydrogels, with one side containing aptamers (control or acrydite), via two-step photo-crosslinking. After fabrication, the bi-phasic samples were incubated in 1 ml of co-culture medium supplemented with 10 ng of VEGF for 1 hr at 37 °C. Following VEGF loading, the hydrogels were washed and incubated with fresh co-culture medium for the study duration. By day 3 of culture, the cells encapsulated within the acrydite aptamer side showed higher cellular structure formation bordering the interface, whereas on the GelMA side cells remained round in morphology suggesting a lower overall cell response (**Figure S6**). In acrydite aptamer functionalized bi-phasic hydrogels, the acrydite aptamer side exhibited an average cell area of 2,569 μm^2^, with 542 μm^2^ for GelMA side. A similar trend was observed for cell aspect ratio. In contrast, for control aptamer functionalized cell-laden bi-phasic hydrogels, both sides of the interface elicit a similar cell responses as indicated by the average cell area (control aptamer side – 3,133 μm^2^ & GelMA side – 2,220 μm^2^) and cell aspect ratio (control aptamer side - 2.8 & GelMA side - 2.9). A lack of response in the control aptamer samples could be explained by the diffusion of aptamer bounded VEGF around the bi-phasic interface, owing to it being only physically entrapped within the polymer network. This local and patterned cellular response confirmed the spatial control over VEGF bioavailability within acrydite aptamer functionalized hydrogels. Interestingly, it is known that matrix bound VEGF facilitates stronger interactions between the VEGF molecules and its receptors presented on endothelial cells, thus leading to an overall improved cellular response.^[24]^ Even though with this aptamer-based system the VEGF sequestration is mechanistically different, the cells showed a superior cell response in acrydite aptamer compared to control aptamer bi-phasic hydrogels **(Figure S6)**. With this logic, the rationale behind the clear advantage of acrydite functionalized aptamers (similar to matrix bound VEGF) over the control aptamers (similar to soluble VEGF) could be understood. Additionally, acrydite aptamers provide a clear benefit of spatial and temporal control over VEGF release.

Neovascularization is an important parameter for the success of engineered tissues, which is highly dependent on the spatiotemporally controlled availability of angiogenic growth factors. For this purpose, we investigated the capacity of bi-phasic hydrogel systems to spatiotemporally control microvascular network organization. To do so, complementary sequence was added into the cell-laden bi-phasic aptamer-functionalized hydrogels on day 4 to trigger VEGF release. The hydrogels treated with or without complementary sequences were compared at different time points (day 5 & 10). Considering the large hydrogel size, we defined four different regions within these sample: (1) immediate vicinity of the interface, marked as near (on both sides) and (2) away from the interface, marked as far (for both sides). It is well established that crosstalk between endothelial cells and MSCs can direct MSC differentiation towards the smooth muscle cell lineage and thus stabilize vascular structures.^[25]^ To investigate this, hydrogels samples were immunostained for both the endothelial cell specific marker von Willebrand factor (vWF), and the smooth muscle marker α-smooth muscle actin (α-SMA).

As evident from confocal microscopy images, all of the acrydite samples display positive staining for vWF and α-SMA at both time points, regardless of complementary sequence addition on day 4 **(Figure 4 & S7)**. The expression of α-SMA and co-localization with endothelial cells indicates hMSCs differentiation towards mural cells within these hydrogels. On analyzing vWF+ vessel networks, it was found that the complementary sequence addition on day 4 significantly affected the developing microvascular networks **(Figure 4A, D & E)**. On day 5, the vessel density in in the different hydrogel regions with no complementary sequence addition was found to be significantly different (Near aptamer - 4.99 & near GelMA - 2.21; p = 0.0421), whereas in the hydrogels with complementary sequence addition the difference was insignificant **(Figure 4D)**. By day 10 of the culture, the difference in vessel density became insignificant across the interface, independent of the complementary sequence addition. However, the difference between samples with and without complementary sequence added increased. The vessel density on near aptamer side of the interface in complementary sequence added hydrogels was 15.94% compared to 4.57% for no complementary sequence hydrogels (p<0.001) **(Figure 4E)**. Similarly, in near GelMA side of the interface by day 10, the hydrogels with complementary sequence showed a vessel density of 12.30% compared to 2.61% for no complementary sequence added hydrogels(p=0.004) **(Figure 4E)**. Similar trends were observed for other vessel network parameters, such as branching density and average vessel length. Interestingly, the average vessel length in the hydrogels with complementary sequence added increased across the interface from day 5 to day 10 in comparison to the hydrogels with no complementary sequence added. On day 5 the average vessel length in hydrogels without complementary sequence was 0.0513 μm (near aptamer side) & 0.012 μm (near GelMA), whereas for hydrogels with complementary sequence it was 0.035 μm (near aptamer) & 0.017 μm (near GelMA). By day 10, the average vessel length increased to only 0.058 μm (near aptamer) & 0.029 μm (near GelMA) for hydrogels with no complementary sequence addition, whereas for hydrogels with complementary sequence addition, it showed 0.211 μm (near aptamer) & 0.145 μm (near GelMA) **(Figure 4D, E)**.

**Figure 4.**
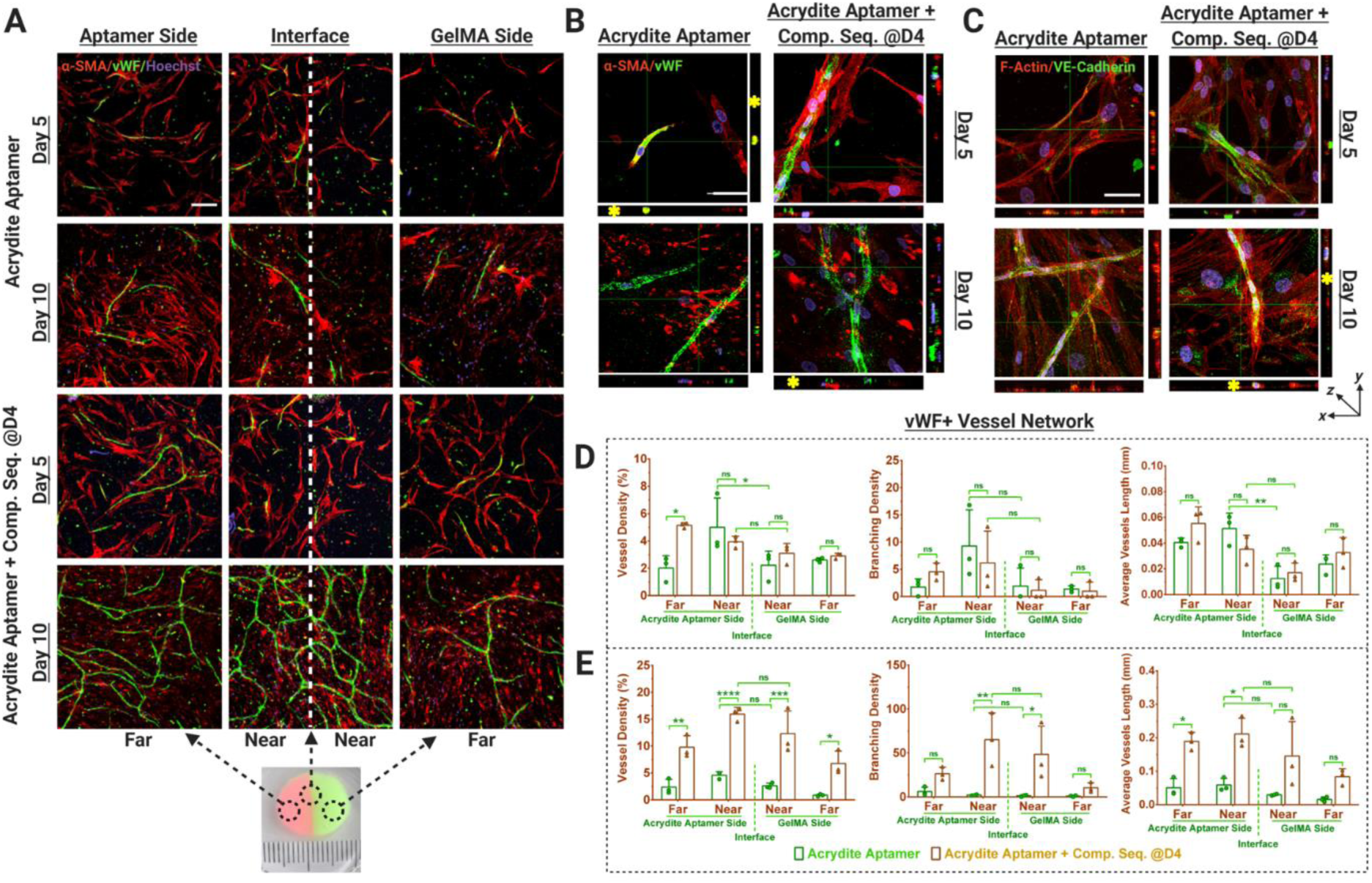
Spatiotemporally controlled vascular network organization within acrydite aptamer functionalized bi-phasic cell-laden hydrogels. (A) Immunostained maximum projection confocal images showing von Willebrand factor (vWF) expression (green) as a marker for endothelial cells and α-smooth muscle actin (α-SMA) (red) as a marker for MSCs differentiation to mural cells, within the HUVECs and hMSCs co-cultured, bi-phasic hydrogels (with or without Comp. Seq. treatment on day 4). The representative photographic image of the actual bi-phasic hydrogel, showing two distinct sides (one with aptamer and other one without) with a distinct interface. Considering its big size, each hydrogel was categorized into four regions; near aptamer- and near GelMA-side being in immediate vicinity of the interface, whereas far aptamer- and far GelMA-side were at the far end from the interface. Scale bar is 200 μm. Orthogonal views of the confocal z-stacks showing (B) vWF + α-SMA + Hoechst stained samples & (C) F-actin Phalloidin + VE-Cadherin + Hoechst stained samples, at higher magnification, showing the developing vascular networks. At the cross-section of the developing vessel, a round lumen like vascular structure could be observed in orthogonal view (indicated by yellow *). The scale bar is 50 μm. Quantification of vWF+ stained vessel network in these samples using Angiotool Software on day 5 (D) and day 10 (E). The values are represented as mean ± SD, along with individual data points. The calculations were performed with three technical replicates, n=3. The statistical significance was calculated using two-way ANOVA with tukey’s post-hoc test where *p<0.05, **p<0.01, ***p<0.001, ****p<0.0001 and ns stands for not significant.

Upon complementary sequence addition in control aptamer-functionalized bi-phasic hydrogels, increased vWF+ vessel network formation was observed on day 5 in comparison to the hydrogels with no complementary sequence addition **(Figure S7A)**. However, no significant difference across the interface was observed, independent of complementary sequence addition **(Figure S7D, E)**. This could be attributed to the diffusion of control aptamer bound VEGF across the interface, as shown previously in aptamer retention **(Figure 1C)** and VEGF ELISA data **(Figure 2C)**. As expected, increased vWF+ network formation was observed in the hydrogels with complementary sequence addition compared to hydrogels with no complementary sequence on day 10 of culture **(Figure S7E)**. As indicated in our previous ELISA results **(Figure 2C)**, complementary sequence addition resulted in VEGF release that in turn homogenized the VEGF availability across the interface, resulting in increased endothelial network formation on day 5. However, hydrogels with no complementary sequence addition showed lower endothelial network formation even after 10 days of culture **(Figure 4A & S7A)**. These observations indicate that VEGF bound to control aptamer does not contribute towards endothelial network formation.

Interestingly, at higher magnifications in orthogonal view, the vWF+ networks showed circular/elliptical lumen like structures wrapped by α-SMA+ cells **(Figure 4B, Figure S7B, Video S1)**. To further asses the vascular stability of the developing microvascular network, the hydrogels were immunostained for expression of the endothelial intercellular junctional protein VE-cadherin at two different time points **(Figure 4C & S7C)**. The positive expression of VE-cadherin in vascular networks confirmed the presence of mature intercellular junctional proteins in all of the samples. As VE-cadherin plays a substantial role in endothelial cell morphogenesis, angiogenesis and vascular stability,^[26]^ the positive expression of VE-cadherin indicates the maturation of these microvascular networks by day 5. As shown in **Figure 4C & S7C**, the VE-cadherin expression in microvascular networks became more evident by day 10 of the culture.

In conclusion, we demonstrated an accessible and flexible method for the spatiotemporal patterning of an angiogenic growth factor in a well-known hydrogel system. This resulted in distinct local differences in vascular network formation in bi-phasic hydrogel systems, where both phases were initially loaded with the same growth factor containing medium. Even though the patterning is simple in this paper, GelMA is known to be compatible with 3D printing and complex photo-patterning techniques, thus extending the possibilities to more complex patterns. Future work will focus on establishing the biological resolution of this technology. Additionally, as aptamer sequences can be designed to capture different proteins, this technology is well adapted for the patterning of different or multiple growth factors. By employing complementary sequences to trigger demanded release, this system has the potential to regulate the availability of multiple growth factors within engineered tissues in space and time.

## Experimental Section

### Materials

Type A 300 bloom porcine skin gelatin (G1890-500G, Sigma Aldrich), Methacrylic anhydride (MA, 276685-500ML, Sigma Aldrich), Dulbecco’s phosphate buffered saline (DPBS, D8537-500ML, Sigma Aldrich), Fisherbrand™ regenerated cellulose dialysis tubing (12-14 kDa, 21-152-14, Fisher Scientific), bovine serum albumin (BSA, A9418, Sigma Aldrich), deuterium oxide (151890, Sigma-Aldrich), 2-hydroxy-4’-(2-hydroxyethoxy)-2-methylpropiophenone (Irgacure 2959, 410896, Sigma Aldrich), VEGF specific control aptamer (47-nt, DNA, IDT), 5’ acrydite modified aptamer (47-nt, DNA, IDT), complementary sequence (Comp. Seq., 46-nt, DNA, IDT), 5’Alexa Fluor 488 modified complementary sequence (Fluoro-Comp. Seq., 46-nt, DNA, IDT), nuclease free water (11-04-02-01, IDT), human VEGF ELISA kit (RAB0507-1KT, Sigma Aldrich), glutaraldehyde solution (340855, Sigma Aldrich), human umbilical vein endothelial cells (HUVECs, C2519A, Lonza), human mesenchymal stem cells (hMSC, PT-2501, Lonza), α –MEM medium (+nucleosides, 22571-020, Gibco), fetal bovine serum (FBS, F7524, Sigma), GlutaMax™ supplement (35050061, Gibco), penicillin-streptomycin (pen/strep, 15140-122, Gibco), L-abscorbic acid (A8960, Sigma Aldrich), trypsin-EDTA 0.25% (+phenol red, 25200072, Gibco), vascular endothelial growth factor 165 human (VEGF, H9166, Sigma Aldrich), Gibco™ FGF-Basic AA 1-155 recombinant human protein (bFGF, PHG0264, Fisher Scientific), endothelial cell basal medium 2 (EGM 2, C-22211, PromoCell), endothelial cell growth medium 2 supplement pack (EGM 2, C-39211, PromoCell), PDMS silicone elastomer (2401673921, Sylgard), live/dead cell double staining kit (04511, Sigma Aldrich), formaldehyde solution (F8775-25ML, Sigma Aldrich), Triton™ X-100 (T8787, Sigma Aldrich), Phalloidin-Atto 488 (49409, Sigma Aldrich), Invitrogen™ Alexa Fluor™ 647 Phalloidin (A22287, Fisher Scientific), Invitrogen™ hoechst 33342 (H1399, Fisher Scientific), monoclonal anti-actin, α-smooth muscle antibody (mouse, A2547, Sigma Aldrich), anti-von Willebrand factor antibody (rabbit, F3520, Sigma Aldrich), human VE-Cadherin antibody (mouse, MAB9381, R&D systems), IgG (H+L) cross-adsorbed goat anti-rabbit, Alexa Fluor® 488, Invitrogen™ (A11008, Fisher Scientific), IgG (H+L) cross-adsorbed goat anti-mouse, Alexa Fluor® 488, Invitrogen™ (A11001, Fisher Scientific), IgG (H+L) highly cross-adsorbed goat anti-mouse, Alexa Fluor® 594, Invitrogen™ (A11032, Fisher Scientific), Corning® 96 well black polystyrene microplate (CLS3603, Sigma), Corning® Costar® ultra-low attachment well plates were used for all cell culture experiments (CLS3473, CLS3474, Sigma).

### Prediction of Secondary Structures

The secondary structures of the DNA based aptamers and their three-dimensional conformations were generated using NUPACK. The presented structure with the lowest free energy at 37 °C was presumed to be the dominant structure **(Figure S1)**.

### Synthesis and characterization of gelatin methacryloyl (GelMA)

Gelatin methacryloyl was synthesized as described previously elsewhere with medium degree of methacryloyl substitution (∼ 60%).^[27]^ Briefly, gelatin was dissolved at 10% (w/v) into DPBS at 60 °C and stirred until fully dissolved. Subsequently, 1.25% (v/v) of MA was added at a rate of 0.5 ml/min to the gelatin solution at 50 °C for 1 hr with continuous stirring. Afterwards, the reaction was stop with additional warm (40 °C) DPBS leading to 5x dilution. The solution was then dialyzed against distilled water using 12-14 kDa cutoff dialysis tubing at 40 °C for 1 week to remove residual salts and methacrylic acid. The solution was freeze dried for 1 week and stored at −20 °C until further use. The degree of functionalization was quantified by H^1^ NMR.^[27]^ The H^1^ NMR spectra were collected at 35 °C in deuterium oxide at a frequency of 400 MHz using NMR spectrometer. The obtained spectral data was analyzed using MestReNova software **(Figure S2)**.

### Synthesis of aptamer-functionalized hydrogel

The freeze-dried GelMA macromer was mixed with Irgacure 2959 as a photoinitiator at 60 °C until fully dissolved. Afterwards, VEGF specific aptamer (Control aptamer or Acrydite aptamer, see **Table S1**) reconstituted in nuclease free water, was added into this solution with 5% (w/v) GelMA, 0.5% (w/v) Irgacure 2959 and 2.5 nmoles of aptamers in every 50 μl of pre-polymer solution. This pre-polymer solution was then used to prepare hydrogel samples for different experiments. As control, GelMA hydrogels were synthesized using same pre-polymer solution but without the aptamer. For photo-crosslinking, requisite amount of pre-polymer solution was added into a PDMS mold, covered with a coverslip and exposed to 0.23 mW/cm^2^ intensity UV light (360-480 nm) (UV-KUB 2, Kloe, France) for 2 min. Samples were detached from the coverslips, transferred to their respective well plates using sterilized spatulas and used for different experiments.

### Evaluation of aptamer retention within the hydrogels

To evaluate aptamer incorporation and their retention within the hydrogels, the aptamer functionalized (Acrydite aptamer or Control aptamer) hydrogels and control GelMA hydrogels were prepared by adding 25 μl of pre-polymer solution into a PDMS mold (2mm thickness and 4mm diameter) following two mins of UV crosslinking. Post photo-crosslinking, the hydrogels were transferred into clear-bottom black polystyrene 96 well plate with one hydrogel per well and were incubated with 2.5 nmoles of Alexa Fluor 488 labelled complementary sequence that binds to the VEGF specific aptamer for 24 hrs in 100 μl DPBS. A mole ratio of 1:1 for aptamer to complementary sequence was maintained. After 24 hrs, the supernatant was discarded and the hydrogels were washed once with DPBS to remove unbound complementary sequence. To this end, 100 μl of fresh DPBS was added into the hydrogel and imaged using fluorescence microscope (EVOS M7000, Thermo Fisher Scientific) as day 1 for the experiment. The experiment was carried out for 10 days where after every 24 hrs, the hydrogel’s supernatant was replaced with fresh 100μl DPBS. The hydrogels were imaged at 60% intensity & 120 ms exposure time; 2x objective. The experiment was performed with three experimental replicates for each group. To quantify the fluorescence intensity, ImageJ software was used.

### Scanning electron microscopy (SEM)

For scanning electron microscopy (SEM), aptamer-functionalized hydrogels with different (acrydite or control) aptamer moles (0.25 nmoles, 2.5 nmoles & 25 nmoles) were prepared as previously discussed. The GelMA hydrogel without any aptamer was prepared as control. For preparing the hydrogel, 500 μl of pre-polymer solution was added into 24 well plate followed by two mins of UV crosslinking. The hydrogel discs of about ∼4 mm thickness and 15 mm diameter were obtained. The hydrogels were washed with DPBS followed by fixing in 2.5% glutaraldehyde for 24 hr at 4 °C. Afterwards, the hydrogels were flash freezed in liquid nitrogen and freeze-dried for 4 days. The freeze-dried hydrogel discs were broken in liquid nitrogen to observe the cross-section, gold-sputtered (Sputter Coater 108 Auto, Cressington Scientific Instruments) and imaged using SEM (JSM-IT100, JEOL). Considering the large hydrogel disc size, three different regions from the same sample were imaged for pore size measurement within each group using ImageJ software.

### Rheological analysis

The viscoelastic properties analysis of the aptamer-functionalized hydrogels with different (acrydite or control) aptamer concentrations (0.25 nmoles, 2.5 nmoles & 25 nmoles) were performed using parallel plate (PP25, 25mm) rheometer (Physica MCR301, Anton Paar). The GelMA hydrogel was prepared as control. Post UV crosslinking, each hydrogel was dispensed onto the rheometer plate (at 20 °C) and the parallel plate was lowered to the desired gap height of 1 mm. The amplitude sweep at a constant 1 rad/s angular frequency (ω) and frequency (f) of 0.159 Hz was carried out over the range of 0.1 till 100 strain (%). The storage module (G’) and loss modulus (G”) were determined as the strain (%) changes.

### Analysis of VEGF sequestration and its triggered release

To conduct VEGF sequestration and triggered release experiments, the aptamer-functionalized hydrogels (2.5 nmoles of acrydite aptamer or control aptamer) were prepared by adding 50 μl of pre-polymer solution into a PDMS mold (2 mm thickness & 6 mm diameter) following two mins of UV crosslinking. Similarly, GelMA hydrogels without aptamers were prepared as controls. Once crosslinked, the hydrogels were transferred to ultra-low attachment 24-well plate. For VEGF loading, the hydrogels were incubated with 1ml of releasing medium (0.1% BSA in DPBS) having 10 ng of VEGF for 1 hr at 37 °C. The mole ratio of aptamer to VEGF was ∼10,000:1. After 1hr incubation, the supernatant was removed and the hydrogels were washed once with releasing medium. Afterwards, the VEGF loaded hydrogels were incubated in 1 ml releasing medium. After every 24 hrs, the supernatant was removed and fresh 1ml releasing medium was added until day 10. The VEGF retention was determined by subtracting the amount of free VEGF in loading solution after 1 hr incubation from the initial loaded VEGF amount. The VEGF loaded aptamer-functionalized hydrogels were also examined for their on-demand triggered release behavior. For this purpose, on day 4, 2.5 nmoles of complementary sequence to VEGF specific aptamer was added to releasing medium making a final volume of 1 ml into “acrydite aptamer + Comp. Seq. @D4 & D9” & “Control aptamer + Comp. Seq, @D4 & D9” samples for 24 hrs. However, in all other hydrogels only releasing medium (no complementary sequence) was added on day 4. Furthermore, on day 9, 2.5 nmoles complementary sequence was added making final volume of 1 ml releasing medium to all aptamer hydrogel samples. The triggered VEGF release was determined by the VEGF amount released on day 5 and 10 (within 24 hrs of complementary sequence addition). All of the supernatants, including loading and washing solutions were stored at −20 C until further analysis.

In order to evaluate the efficacy of ELISA assay in identifying VEGF molecules bounded to control aptamer in supernatant solution, separate experiment was designed. For this purpose, 2.5 nmoles control aptamer and 10 ng VEGF was directly added into 1 ml releasing medium and allowed to incubate for 24 hrs at 37 °C. As a control, only VEGF was added into 1ml releasing medium. The ultra-low attachment 24-well plate was used for this experiment to avoid protein adsorption by the well plate. After 24 hrs, the solution was collected and stored at −20 °C. The amount of VEGF in supernatants was measured by VEGF ELISA kit as per manufacturer’s instructions. The absorbance for the samples were measured using microplate reader (Infinite® 200PRO, Tecan) at 405 nm. Prior to analysis, supernatants were diluted with the sample diluent to ensure the VEGF concentrations within the detectable range of the assay. The absorbance was referenced by subtracting it from the absorbance of zero VEGF concentration. The ELISA data was analysed using GraphPad Prism software. The experiment was performed with three experimental replicates.

### Cell culture

The human umbilical vein derived endothelial cells (HUVECs, Lonza) and human mesenchymal stromal cells (hMSCs, Lonza) were cultured as per the standard protocol. In brief, HUVECs were cultured in EGM-2 medium (EGM-2 Basal medium + EGM-2 Supplements) with 1% (v/v) Pen/Strep. However, for hMSCs α-MEM medium supplemented with 10% (v/v) FBS, 2 mM L-glutamine, 0.2 mM ascorbic acid, 1 % (v/v) Pen/Strep and 1 ng/ml bFGF was used. Both cell types were cultured in a humidified atmosphere with 5% CO2 at 37 °C and passaged at 80% confluence.

### 3D Culture in aptamer-functionalized hydrogels

For co-culture experiments, 1:1 ratio of HUVECs to hMSCs with a total seeding density of 2.5 x 10^6^ cells/ml was used throughout this study. Similarly, hMSCs medium (α-MEM, 10% FBS, 2 mM L-glutamine, 0.2 mM abscorbic acid & 1% Pen/Strep) and HUVECs medium (EGM-2 Basal medium + 1% Pen/Strep + all EGM-2 supplements except for VEGF (C-30260, PromoCell)) were used in 1:1 ratio as co-culture medium without VEGF supplement. In all experiments, both cell types between passage 3 and 5 were used.

For 3D culture experiments, HUVECs (P3) and hMSCs (P4) were trypsinized, counted (using trphan blue) and were re-suspended in 1:1 ratio with total seeding density of 2.5 x 10^6^ cells/ml. This cell suspension was further centrifuged at 300 g for 3 mins at 4 °C to get the cell pellet. The supernatant was removed and pre-polymer solution (5% GelMA + 0.5% Irgacure + 2.5 nmoles acrydite or control aptamer in DPBS) was directly added and mixed gently by pipetting. Around 50 μl from this pre-polymer solution with cells was dispensed into ultra-low attachment 96 well-plate, followed by 2 mins of UV crosslinking. The experiment was performed under aseptic conditions. Afterwards, 200 μl of co-culture medium along with 10 ng VEGF was added onto the hydrogels and were kept inside humidified incubator at 37 °C with 5% CO2 for 1 hr. After 1 hr of VEGF loading, supernatant was removed, hydrogels were washed twice with DPBS and then 200 μl of co-culture medium was added. The medium was changed after every 24 hrs throughout the study duration. As a control, GelMA hydrogel with cells encapsulated in it were also prepared similarly without the aptamer. To observe the effect of triggered release of VEGF on the cells, 2.5 nmoles of complementary sequence (with aptamer to complementary sequence ratio of 1:1) was added onto the 200 μl of co-culture medium on day 4. These hydrogels were analyzed for cell viability on day 1 and day 5 by using Live/Dead cell viability kit as per manufacturer’s protocol. Briefly, the culture medium of the hydrogel was removed and 100 μl of staining solution (2 μl of solution A + 1 μl of solution B in 1 ml of DPBS) was added, following 15 min incubation. The stained hydrogels were imaged using fluorescent microscope (EVOS M7000, Thermo Fisher Scientific). For this study, three experimental replicates were used. The live (green) and dead (red) stained cells were counted using ImageJ software using atleast 5 images per hydrogel.

### Bi-phasic cell-laden hydrogels

To fabricate bi-phasic cell-laden hydrogels, autoclaved PDMS molds (3 mm thickness & 15 mm diameter) with its removable halves insert were used. To start with, HUVECs (P3) and hMSCs (P4) were trypsinized, counted (using tryphan blue) and were re-suspended in 1:1 ratio with final seeding density of 2.5 x 10^6^ cells/ml. To prepare two different pre-polymer solutions, the cell suspension with same seeding density was centrifuged at 300 g for 3 mins at 4 °C in two separate 15 ml falcon tubes. For pre-polymer solution 1, supernatant was removed and pre-polymer solution (5% GelMA + 0.5% Irgacure 2959 + 2.5 nmoles acrydite or control aptamer in DPBS) was added and mixed gently by pipetting. Similarly, for pre-polymer solution 2, supernatant was removed and 5% GelMA + 0.5% Irgacure 2959 in DPBS solution was added. Afterwards, PDMS molds with one insert covering its half area were placed in sterile cell culture petri dish (90 mm diameter) and 200 μl of pre-polymer solution 1 was added. The mold was covered with coverslip and exposed for UV crosslinking (1 min, 0.23 mW/cm^2^ intensity). After the crosslinking of one side, the PDMS mold insert and coverslip was carefully removed; and pre-polymer solution 2 was added in the rest of the mold followed by UV crosslinking (1min, 0.23 mW/cm^2^ intensity). Thereafter, the coverslip and PDMS molds were removed and bi-phasic cell-laden hydrogels were transferred to ultra-low attachment 24-well plate using sterilized spatula. Herein, for VEGF loading, 1 ml of co-culture medium with 10 ng VEGF was added into each sample and incubated at 37 °C for 1 hr. Subsequently, the medium was replaced with fresh 1ml co-culture medium. The medium was changed once in everyday throughout the experiment. To evaluate the effect of triggered VEGF release on bi-phasic cell-laden hydrogels, 2.5 nmoles complementary sequence (1:1 mole ratio of aptamer to complementary sequence) into 1ml co-culture medium was added for acrydite and control aptamer based bi-phasic hydrogels on day 4 for 24 hrs. These samples were analyzed at different time-points.

### Immunostainings

In prior to staining, all of the samples were fixed using 4% formaldehyde solution in DPBS for 30 mins, washed with DPBS and permeabilized using 0.1% Triton X-100 in DPBS. Afterwards, the samples were washed with DPBS and blocking solution of 1% FBS in DPBS was added for 45 mins. For actin cytoskeleton staining of bi-phasic cell-laden hydrogels, the samples were incubated with Phalloidin - Atto 488 (1:50 in DPBS) or Alexa Fluor 647 Phalloidin (1:40 in DPBS) - for 1 hr at room temperature, followed by 10 mins incubation with Hoechst 33342 (1:2000 in DPBS). For immunostainings, post blocking solution, the samples were washed with DPBS and incubated for overnight at 4 °C with monoclonal anti-actin, α-smooth muscle antibody (mouse, 1:300 in DPBS) and anti-von Willebrand factor antibody (rabbit, 1:200 in DPBS). The samples were then washed with DPBS, IgG (H+L) highly cross-adsorbed goat anti-mouse, Alexa Fluor® 594 (1:1000 in DPBS) and IgG (H+L) cross-adsorbed goat anti-rabbit, Alexa Fluor® 488 (1:1000 in DPBS) solutions were added for 2 hrs in dark. Subsequently, the samples were washed with DPBS and incubated with Hoechst 33342 (1:2000) for 10 mins in dark. For tight-junctions specific proteins staining, after blocking solution & DPBS washing, the samples were incubated for overnight at 4 °C with human VE-Cadherin antibody (mouse, 1:200 in DPBS). The samples were washed with DPBS and incubated with IgG (H+L) cross-adsorbed goat anti-mouse, Alexa Fluor® 488 (1:1000 in DPBS) for 2 hrs in dark. After 1 hr, Phalloidin 647 (1:40 in DPBS) was added into the secondary antibody staining solution and allowed to incubate for 1 hrs in dark. Thereafter, the samples were washed and Hoechst 33342 (1:2000) was added for 10 mins in dark. Once stained, samples were washed with DPBS, 500 μl DPBS was added and stored at 4 °C until imaged.

### Statistical Analysis

Statistical analysis was performed using GraphPad Prism 7 software. Two-way analysis of variance (ANOVA) with Tukey’s multiple comparisons test was used to analyze the data. Significance was set at p<0.05. The data is represented as mean ± SD.

## Supporting Information

Supporting Information is available.

## Acknowledgements

This work was supported by the European Research Council (ERC) under the European Union’s Horizon 2020 Research and Innovation Programme (No. 724469). Illustrations for the manuscript were created with BioRender.com.

## References

[1] J. Rouwkema, A. Khademhosseini, Trends Biotechnol. 2016, 34, 733.

[2] K. M. Park, S. Gerecht, Development 2014, 141, 2760.

[3] M. M. Martino, S. Brkic, E. Bovo, M. Burger, D. J. Schaefer, T. Wolff, L. Gürke, P. S. Briquez, H. M. Larsson, R. Gianni-Barrera, J. A. Hubbell, A. Banfi, Front. Bioeng. Biotechnol. 2015, 3, 45.

[4] N. Ferrara, Mol. Biol. Cell 2010, 21, 687.

[5] P. Carmeliet, Nat. Med. 2003, 9, 653.

[6] A. A. Ucuzian, A. A. Gassman, A. T. East, H. P. Greisler, J. Burn Care Res. 2010, 31, 158.

[7] P. M. Gawade, J. A. Shadish, B. A. Badeau, C. A. DeForest, Adv. Mater. 2019, 31, 1902462.

[8] A. M. Rosales, K. S. Anseth, Nat. Rev. Mater. 2016, 1, 15012.

[9] E. R. Ruskowitz, C. A. Deforest, Nat. Rev. Mater. 2018, 3, 17087.

[10] C. A. Deforest, D. A. Tirrell, Nat. Mater. 2015, 14, 523.

[11] T. Kamperman, M. Koerselman, C. Kelder, J. Hendriks, J. F. Crispim, X. de Peuter, P. J. Dijkstra, M. Karperien, J. Leijten, Nat. Commun. 2019, 10, 4347.

[12] C. Fan, J. Shi, Y. Zhuang, L. Zhang, L. Huang, W. Yang, B. Chen, Y. Chen, Z. Xiao, H. Shen, Y. Zhao, J. Dai, Adv. Mater. 2019, 31, 1902900.

[13] J. R. García, A. Y. Clark, A. J. García, J. Biomed. Mater. Res. - Part A 2016, 104, 889.

[14] J. Ishihara, A. Ishihara, K. Fukunaga, K. Sasaki, M. J. V. White, P. S. Briquez, J. A. Hubbell, Nat. Commun. 2018, 9, 2163.

[15] S. Prakash Parthiban, D. Rana, E. Jabbari, N. Benkirane-Jessel, M. Ramalingam, Acta Biomater. 2017, 51, 330.

[16] G. A. Hudalla, W. L. Murphy, Adv. Funct. Mater. 2011, 21, 1754.

[17] A. S. R. Potty, K. Kourentzi, H. Fang, G. W. Jackson, X. Zhang, G. B. Legge, R. C. Willson, Biopolymers 2009, 91, 145.

[18] M. Kimoto, R. Yamashige, K. I. Matsunaga, S. Yokoyama, I. Hirao, Nat. Biotechnol. 2013, 31, 453.

[19] M. R. Battig, B. Soontornworajit, Y. Wang, J. Am. Chem. Soc. 2012, 134, 12410.

[20] T. Buie, J. McCune, E. Cosgriff-Hernandez, Trends Biotechnol. 2020, 38, 546.

[21] B. Soontornworajit, J. Zhou, M. T. Shaw, T. H. Fan, Y. Wang, Chem. Commun. 2010, 46, 1857.

[22] N. Zhao, A. Suzuki, X. Zhang, P. Shi, L. Abune, J. Coyne, H. Jia, N. Xiong, G. Zhang, Y. Wang, ACS Appl. Mater. Interfaces 2019, 11, 18123.

[23] L. Abune, N. Zhao, J. Lai, B. Peterson, S. Szczesny, Y. Wang, ACS Biomater. Sci. Eng. 2019, 5, 2382.

[24] T. T. Chen, A. Luque, S. Lee, S. M. Anderson, T. Segura, M. L. Iruela-Arispe, J. Cell Biol. 2010, 188, 595.

[25] T. P. Lozito, C. K. Kuo, J. M. Taboas, R. S. Tuan, J. Cell. Biochem. 2009, 107, 714.

[26] D. Vestweber, Arterioscler. Thromb. Vasc. Biol. 2008, 28, 223.

[27] J. W. Nichol, S. T. Koshy, H. Bae, C. M. Hwang, S. Yamanlar, A. Khademhosseini, Biomaterials 2010, 31, 5536.

