## Supplemental information for "Aptamer based spatiotemporally controlled growth factor patterning for tunable local microvascular network formation in engineered tissues"

#### Supporting Information

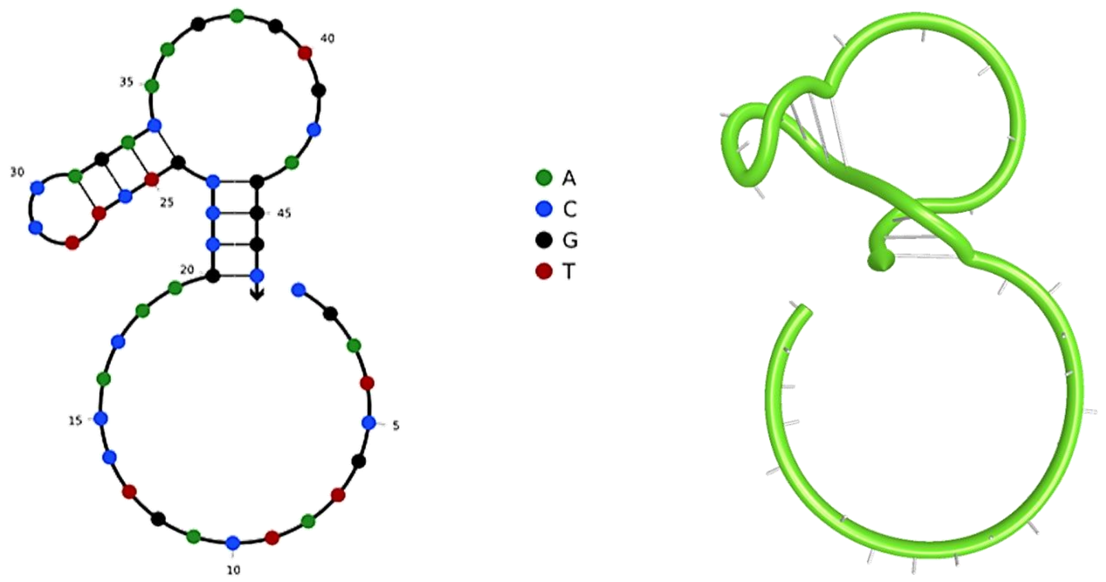


**Figure S1.** Predicted secondary structures and three-dimensional configuration of the VEGF specific aptamer used in this study. The secondary structures of the DNA based aptamers and their three- dimensional conformations were generated using NUPACK software.

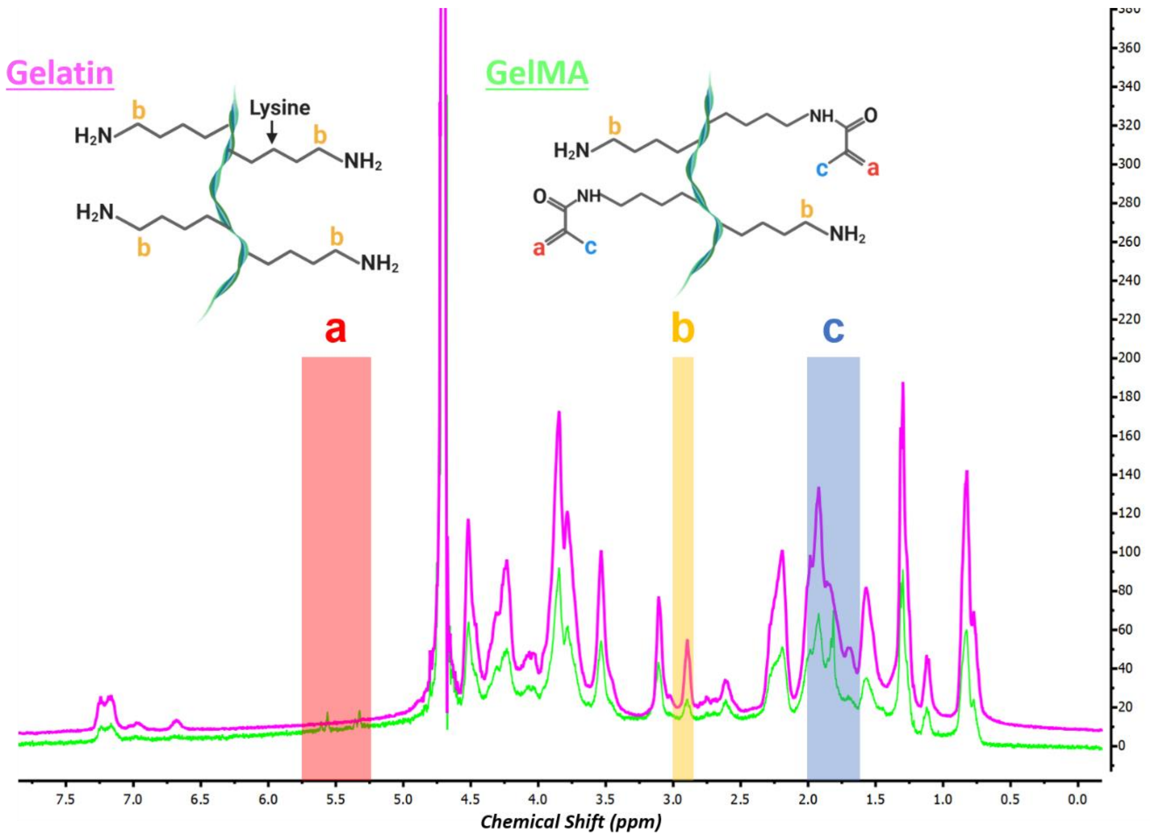


**Figure S2.** NMR Spectrum of gelatin methacryloyl (GelMA) and gelatin. The highlighted red "a" represents the signals of methyl group and yellow "b" shows the acrylic protons of the grafted methacrylic group; and blue "c" indicates the signal of lysine methylene.

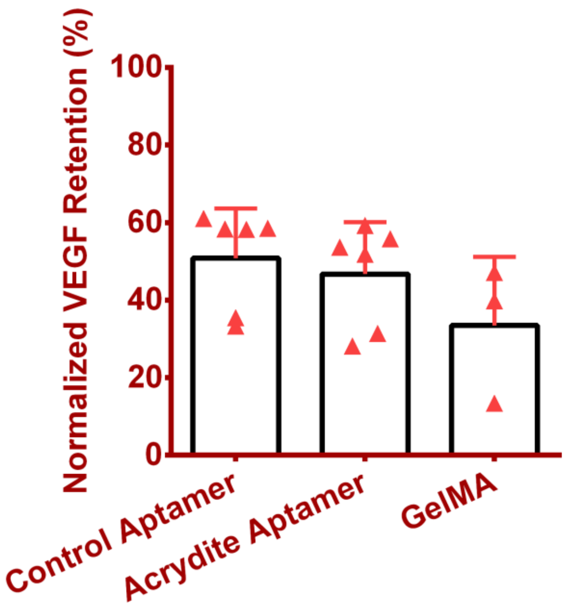


**Figure S3.** The normalized VEGF retention % after 1 hr incubation with the control aptamer-, acrydite aptamer-functionalized and GelMA hydrogel samples. The aptamer concentration within hydrogels was fixed at 2.5 nmoles and loaded with 10 ng VEGF in 1ml loading solution. The data is normalized with control PBS samples (data not shown). The quantification was performed using ELISA assay having n=6 (for aptamer samples) and n=3 (for GelMA), experimental replicates. The data is represented as mean ± S.D. with individual data points.

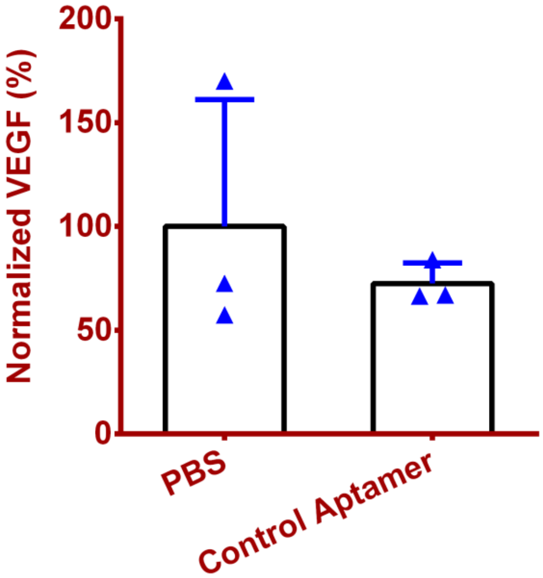


**Figure S4.** Normalized VEGF % in presence of control aptamer (2.5 nmoles) and only PBS after 24 hr incubation at 37 °C. The data is normalized with PBS samples. The graph indicates the difference in ELISA sensitivity in VEGF detection in the PBS versus in presence of control aptamers, where (part of) the VEGF will be in a bound state with the aptamer. The quantification was performed using ELISA assay with n=3 experimental replicates. The data is represented as mean ± S.D. with individual data points.

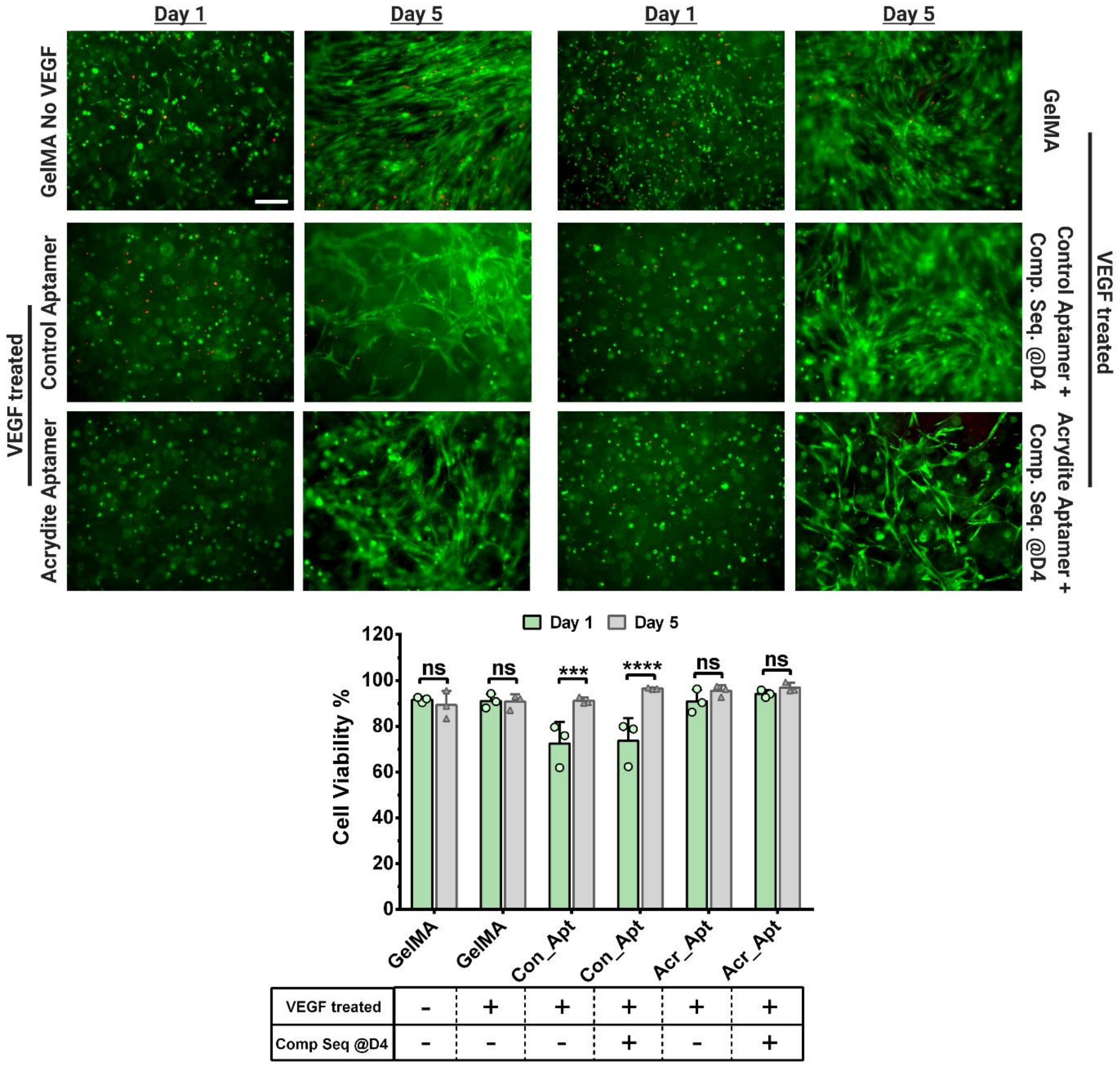


**Figure S5.** Cell viability of the 3D cultured aptamer functionalized hydrogels with or without VEGF loading. Live/Dead stained fluorescent microscopic images of 3D cultured hydrogels with control/acrydite aptamer functionalized hydrogels in the presence or absence of Comp. Seq. addition on day 4, on different time points. The HUVECs and hMSCs were co-cultured in 1:1 ratio. For VEGF loading, the hydrogels were incubated with co-culture medium (without VEGF supplement) supplemented with 10 ng external VEGF, for 1 hr at 37 °C. As a control, GelMA hydrogels, with or without 1 hr VEGF loading were also studied. The scale bar is 200 µm. Cell viability % quantification was performed using ImageJ software. The quantification was performed with three experimental replicates, n=3. The statistical significance was calculated using two-way ANOVA with Bonferroni's multiple comparisons test where ***p=0.0005, ****p<0.0001 and ns stands for not significant.

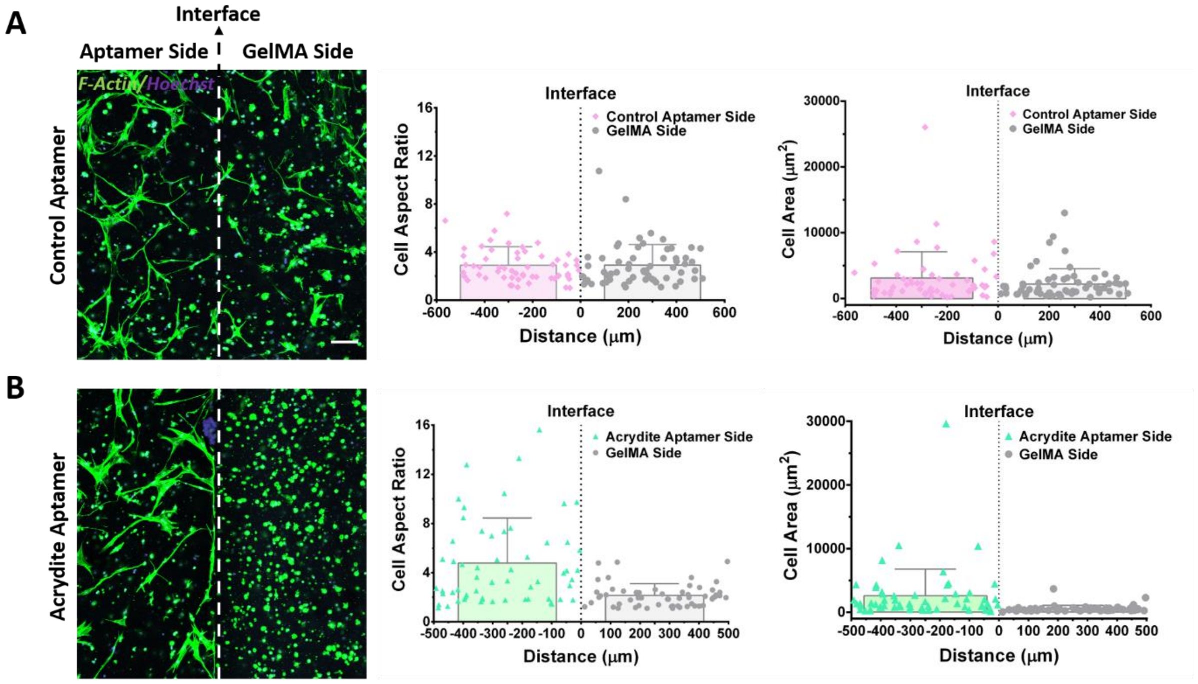


**Figure S6.** Cellular response within bi-phasic cell-laden hydrogels on day 3. The maximum projection confocal images of the HUVECs and hMSCs, co-cultured within $VEGF_{165}$ loaded (A) control aptamer- and (B) acrydite aptamer-functionalized bi-phasic hydrogels. For the ease of understanding, each bi-phasic hydrogel was categorized into two regions; aptamer side and GelMA side where the white dashed line indicates the interface. The cells were stained with cytoskeletal F-actin Phalloidin (green) and Hoechst (blue). The scale bar is 100 µm. Quantification of the individual cell area plotted against the distance from the interface within the (A) control aptamer- and (B) acrydite aptamer-functionalized bi-phasic hydrogels. The quantification was performed using ImageJ software, where values are represented in scatter plot with individual data points along with overall mean ± S.D. The calculations were performed with three technical replicates, n=3.

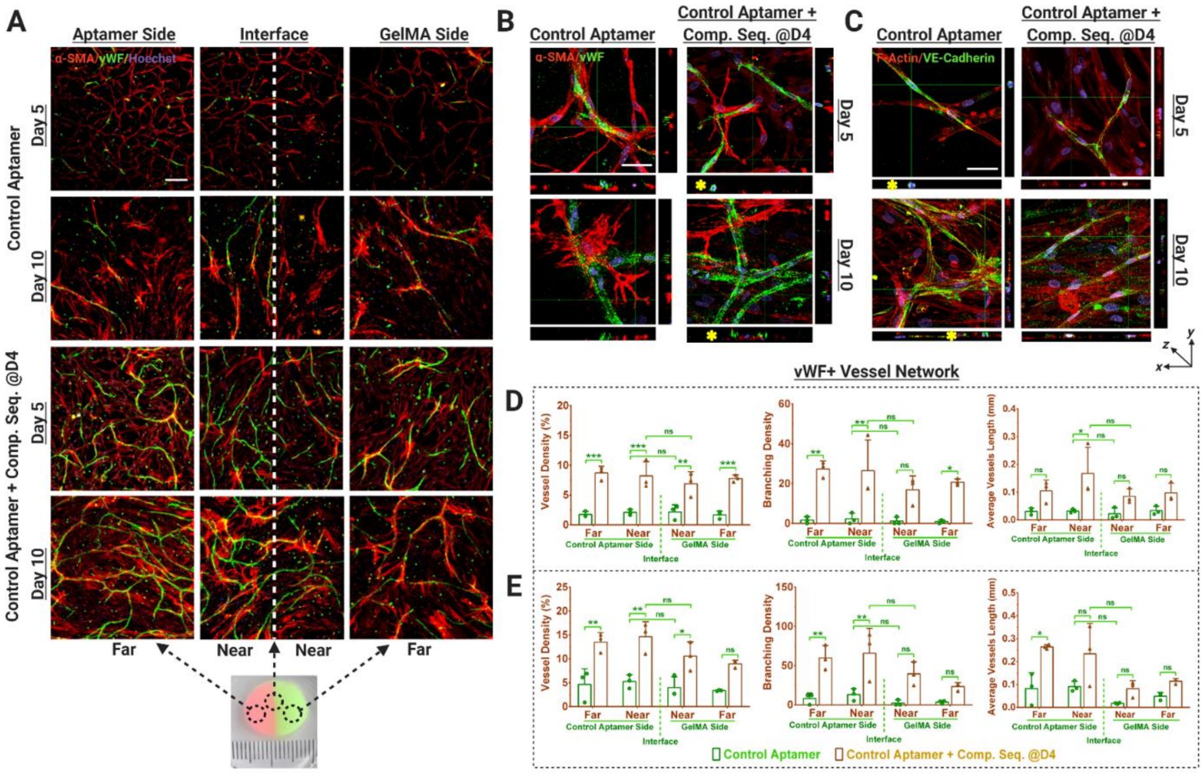

**Figure S7.** Vascular network organization within control aptamer functionalized bi-phasic cell-laden hydrogels. (A) Immunostained maximum projection confocal images showing von Willebrand factor (vWF) expression (green) as a marker for endothelial cells and α-smooth muscle actin (α-SMA) (red) as a marker for MSCs differentiation to mural cells, within the HUVECs and hMSCs co-cultured, bi-phasic hydrogels (with or without complementary sequence treatment on day 4). The representative photographic image of the actual bi-phasic hydrogel, showing two distinct sides (one with aptamer and other one without) with a distinct interface. Considering its big size, each hydrogel was categorized into four regions; near aptamer- and near GelMA-side being in immediate vicinity of the interface, whereas far aptamer- and far GelMA-side were at the far end from the interface. Scale bar is 200 µm. Orthogonal views of the confocal z-stacks showing (B) vWF + α-SMA stained samples & (C) F-actin Phalloidin + VE-Cadherin stained samples, at higher magnification, showing the developing vascular networks. At the cross-section of the developing vessel, a round lumen like vascular structure could be observed in orthogonal view. The scale bar is 50 µm. Quantification of vWF+ stained vessel network in these samples using Angiotool Software on day 5 (D) and day 10 (E). The values are represented as mean ± SD, along with individual data points. The calculations were performed with three technical replicates, n=3. The statistical significance was calculated using two-way ANOVA with tukey's post-hoc test were *p<0.05, **p<0.01, ***p<0.001, ****p<0.0001 and ns means not significant.

### SUPPORTING TABLES & VIDEO

**Table S1.** Full sequences and other characteristics of the aptamers used in this study. Tm denotes the melting temperature (50 mM NaCl), MW is molecular weight and N signifies the number of nucleotides.

| Aptamer | Sequence (5’→ 3’) | Tm | MW | N |
|---|---|---|---|---|
| **Control Aptamer** | CGA TCG TAT CAG TCC ACA AGC CCG TCT TCC AGA CAA GAG TGC AGG GC | 70.8 °C | 14418.4 | 47 |
| **Acrydite Aptamer** | /5Acryd/CGA TCG TAT CAG TCC ACA AGC CCG TCT TCC AGA CAA GAG TGC AGG GC | 70.8 °C | 14665.6 | 47 |
| **Comp. Seq.** | CGC CCT GCA CTC TTG TCT GGA AGA CGG GCT TGT GGA CTG ATA CGA TCG | 71.3 °C | 14791.6 | 48 |
| **Fluoro – Comp. Seq.** | /5Alexa488N/CGC CCT GCA CTC TTG TCT GGA AGA CGG GCT TGT GGA CTG ATA CGA TCG | 71.3 °C | 15487.2 | 48 |

**Video S1.** Immunostained von Willebrand factor (vWF) in green color (endothelial cells marker) and α-smooth muscle actin (α-SMA) in red color indicating MSCs differentiation to mural cells, within the HUVECs and hMSCs co-cultured, acrydite aptamer-functionalized bi-phasic hydrogel showing smooth muscle-like cells wrapping around the developing endothelial network for support.
